# The Cardiac Calcium Handling Machinery is Remodeled in Friedreich’s Ataxia

**DOI:** 10.1101/2023.11.09.566141

**Authors:** Roman Czornobil, Obada Abou-Assali, Elizabeth Remily-Wood, David R Lynch, Sami F. Noujaim, Bojjibabu Chidipi

## Abstract

**Background:** Friedreich’s ataxia (FA) is an inherited neurodegenerative disorder that causes progressive nervous system damage resulting in impaired muscle coordination. FA is the most common autosomal recessive form of ataxia and is caused by an expansion of the DNA triplet guanine–adenine–adenine (GAA) in the first intron of the Frataxin gene (FXN), located on chromosome 9q13. In the unaffected population, the number of GAA repeats ranges from 6 to 27 repetitions. In FA patients, GAA repeat expansions range from 44 to 1,700 repeats which decreases frataxin protein expression. Frataxin is a mitochondrial protein essential for various cellular functions, including iron metabolism. Reduced frataxin expression is thought to negatively affect mitochondrial iron metabolism, leading to increased oxidative damage. Although FA is considered a neurodegenerative disorder, FA patients display heart disease that includes hypertrophy, heart failure, arrhythmias, conduction abnormalities, and cardiac fibrosis.

**Objective:** In this work, we investigated whether abnormal Ca^2+^ handling machinery is the molecular mechanism that perpetuates cardiac dysfunction in FA.

**Methods:** We used the frataxin knock-out (FXN-KO) mouse model of FA as well as human heart samples from donors with FA and from unaffected donors. ECG and echocardiography were used to assess cardiac function in the mice. Expression of calcium handling machinery proteins was assessed with proteomics and western blot. In left ventricular myocytes from FXN-KO and FXN-WT mice, the IonOptix system was used for calcium imaging, the seahorse assay was utilized to measure oxygen consumption rate (OCR), and confocal imaging was used to quantify the mitochondrial membrane potential (Δψm) and reactive oxygen species (ROS).

**Results:** We found that major contractile proteins, including SERCA2a and Ryr2, were downregulated in human left ventricular samples from deceased donors with FA compared to unaffected donors, similar to the downregulation of these proteins in the left ventricular tissue from FXN-KO compared to FXN-WT. On the ECG, the RR, PR, QRS, and QTc were significantly longer in the FXN-KO mice compared to FXN-WT. The ejection fraction and fractional shortening were significantly decreased and left ventricular wall thickness and diameter were significantly increased in the FXN-KO mice versus FXN-WT. The mitochondrial membrane potential Δψm was depolarized, ROS levels were elevated, and OCR was decreased in ventricular myocytes from FXN-KO versus FXN-WT.

**Conclusion:** The development of left ventricular contractile dysfunction in FA is associated with reduced expression of calcium handling proteins and mitochondrial dysfunction.

## 1. INTRODUCTION

FA is a genetic neurodegenerative disease. FA affects almost 1 in 50,000 people with equal frequency in both sexes and is typically diagnosed in mid-childhood^1^. FA is the most common autosomal recessive form of ataxia and is caused by an expansion of the DNA triplet guanine–adenine–adenine (GAA) in the first intron of the Frataxin gene (FXN), located on chromosome 9q13^2^. In the unaffected population, the number of GAA repeats generally ranges from 6 to 27 repetitions, but in FA patients, GAA repeat expansions range from 44 to 1,700 repeats^3^. This high number of GAA repeats decreases frataxin expression in FA patients. Frataxin is a 210 amino acid protein, ubiquitously expressed in the whole body, with high levels found in the heart, spinal cord, liver, pancreas, and muscle^4^. Frataxin plays a significant role in iron metabolism, iron storage, and iron-sulfur cluster biosynthesis in the mitochondria^5^. It is thought that decreased expression of frataxin, as in FA, impairs mitochondrial function by impairing the iron-sulfur cluster, therefore affecting heme biosynthesis and ultimately leading to increased oxidative stress^1 6, 7^ ^14^. Decreased limbs and trunk movements, dysarthria, and heart disorders are clinical symptoms of FA ^8^.

In FA, the leading causes of mortality are cardiac dysfunction (59%), non-cardiac abnormalities (27.9%), and unknown (9.8%) causes^9^. Hypertrophic cardiomyopathy (HCM) is the most commonly occurring cardiac disease in FA^10–12^. Left ventricle (LV) wall thickness, LV mass, and LV ejection fraction are significantly changed in patients with FA HCM^10^. Additionally, atrial tachycardia^13^, and intra atrial block are also found in FA patients^14^. The interatrial block most likely leads to the increased P-wave duration that is manifested in FA^15^. Norrish G et al. conducted a study on a large cohort of FA patients and found that heart failure is a major cause of death in this patient population^12^. Additionally, the development of left ventricular systolic dysfunction has been shown to lead to a poorer prognosis in FA patients^16–18^. Despite the strong cardiac phenotype associated with FA, there is no solid understanding of the progression of cardiac abnormalities in FA.

Ca^2+^ plays a crucial role in the regulation of cellular function as a major intracellular messenger. Intracellular changes in Ca^2+^ concentration regulate cardiac contractility. During systole, Ca^2+^ ions enter the myocyte through L-type Ca^2+^ channels located mainly in the transverse tubules. This leads to the release of Ca^2+^ from the sarcoplasmic reticulum (SR) through the type 2 ryanodine receptors (Ryr2) in a process known as Ca^2+^ induced Ca^2+^ release. The free Ca^2+^ then binds to troponin C which allows for the cross-bridge formation of contractile machinery and leads to muscle contraction. This free Ca^2+^ can also be taken into the mitochondria for the generation of ATP via activation of cellular processes, such as the citric acid cycle and oxidative phosphorylation. Ca^2+^ transport to and from the mitochondria is tightly regulated by several mitochondrial membrane proteins, including voltage-dependent anion channel (VDAC), mitochondrial Ca^2+^ uniport (MCU), and the mitochondrial sodium-Ca^2+^ exchanger (NCLX). During diastole, Ca^2+^ is then removed from the cytoplasm for relaxation to occur, where Ca^2+^ is pumped back into the SR by the SERCA2a and extruded out of the cell through the sodium-Ca^2+^ exchanger (NCX). Alterations or mutations in Ca^2+^ handling proteins, such as Ryr2, SERCA2a, and NCX can result in cardiac pathology and arrhythmias^19–21^. Additionally, irregularities in mitochondrial machinery can be detrimental to cardiomyocyte survival, resulting in progression to heart failure^22–24^.

The correlation between cardiac involvement in FA and the decrease of frataxin affecting cardiac contractility has not been analyzed. In this study, we investigated whether the Ca^2+^ handling machinery is remodeled in FA, and we found that Ca^2+^ handling was dysregulated, and cardiac contractility and cellular function were decreased.

## 2. METHODS

### 2.1 FA animal model

We used the FA mouse model B6.Cg-Fxn^em2Lutzy^ Fxn^em2.1Lutzy^Tg(Ckmm-cre)5Khn/J (FXN-KO) and littermates control Heterozygous for Fxn^em2Lutzy^, Wildtype for Fxn^em2.1Lutzy^, Hemizygous for Tg(Ckmm-cre)5Khn (FXN-WT) purchased from the Jackson Laboratory (Stock# 029720)^25^. We also generated these FXN-KO and FXN-WT mice (Stock# 029720) by breeding male C57BL/6J-congenic *Fxn^em2Lutzy^* (homozygous frataxin floxed exon 2) (Stock# 028520) and female C57BL/6-congenic *Fxn^em^*^2^*^.1Lutzy^* Tg(Ckmm-cre)5Khn (heterozygous frataxin global knockout and homozygous MCK-Cre) (Stock# 029100), and these breeders were purchased from the Jackson Laboratory ^26^. For euthanasia, CO_2_ was used to induce loss of consciousness, and after the loss of the pinch reflex, swift cervical dislocation was performed, followed by rapid harvesting of the heart via thoracotomy. This method is consistent with the guidelines of the American Veterinary Medical Association.

### 2.2 Human ventricular tissues

We obtained left ventricular samples from five different decedent donors with FA from Friedreich’s Ataxia Research Alliance (FARA) tissue bank at Albany VA Hospital. Left ventricular samples were also obtained from five age, and sex matched samples from donors with no diagnosis of FA. These control hearts were rejected for transplant and were obtained through a USF agreement with Lifelink®, an FDA-approved, Tampa based organization for organ procurement (listed in Table 1).

**Table 1.** Demographics of left ventricular human samples used to perform western blotting experiments from the control unaffected individuals and FA patient’s hearts.

| **Name** | **Gender** | **Age** |
| --- | --- | --- |
| <u>Control hearts</u> |  |  |
| 1 | M | 68 |
| 2 | M | 36 |
| 3 | F | 68 |
| 4 | F | 29 |
| 5 | F | 30 |
| <u>FA patient's hearts</u> |  |  |
| 1 | M | 80 |
| 2 | M | 35 |
| 3 | F | 67 |
| 4 | F | 28 |
| 5 | F | 26 |

### 2.3 ECG

Mice were anesthetized with 1.8% isoflurane. The ECG was recorded in Lead II configuration via an Animal Bio Amp (AD Instruments, USA) and digitized with the PowerLab data acquisition system (AD Instruments, USA). LabChart Pro 7.2 software (AD Instruments, USA) was used for the acquisition and analysis of cardiac electrical signals.

### 2.4 Echocardiography

Transthoracic echocardiography was performed using a Vevo3100LT Ultrasonograph (Visual Sonics) equipped with a 30 MHz transducer. Cardiac function was measured in both FXN-WT and FXN-KO mice. Briefly, the mice were anesthetized with 1–1.5% isoflurane, and body temperature was maintained at 37 °C. First, a 2-D mode in the parasternal long axis was imaged to identify the left ventricle part and then switched to the parasternal short axis at the mid-papillary muscle to image the thickness of the wall. From this parasternal short-axis view, the 2-D guided M-mode across the anterior wall and posterior wall were recorded. Left ventricular anterior wall (LVAW), left ventricular interior diameter (LVID), and left ventricular posterior wall thickness (LVPW) at systole and diastole were measured. Ejection fraction (EF) % was calculated as (EDV − ESV)/EDV*100%, and fractional shortening (FS) % was calculated by (LVID;d − LVID;s)/LVID;d*100%.

### 2.5 Western blotting

For collecting lysates from the left ventricle, samples from FXN-KO, FXN-WT, human FA and control ventricular tissues were flash frozen in liquid nitrogen, and tissues were cryopowdered with a Bessman tissue pulverizer (Spectrum Med, San Diego, CA, USA). Protein isolation was completed by adding 10 µL/mg of a RIPA lysis buffer to the cryopowder. The total lysate was centrifuged at 12,000 rpm for 20 min, and the supernatant was collected. Protein concentrations were determined with a BCA assay (Pierce™ BCA Protein Assay Kit; #23225, ThermoFisher, Carlsbad, CA, USA). Total protein (5∼15 µg) was loaded and separated by 4–20% pre-made SDS-PAGE gels (#4568095, BioRad, Mini-PROTEAN TGX Stain-Free Precast Gels, Irvine, CA, USA), then transferred to a nitrocellulose membrane, and probed with the following primary antibodies: mouse monoclonal anti-Frataxin (dilution 5 µg/ml; Abcam; Catalog # ab110328), and mouse monoclonal anti-Ryr2 (dilution 1:1000; Santa Cruz Biotechnology; Catalog # sc-376507), mouse monoclonal anti-SERCA2a (dilution 1:500; Santa Cruz Biotechnology; Catalog # sc-376235). The following secondary antibodies were used: anti-mouse IgGk BP-HRP (dilution 1:500; Santa Cruz Biotechnology; Catalog # sc-516102) for detecting Ryr2, fluorescent IRDye 680 goat anti-mouse antibodies (1:25,000; LiCOR; Catalog # 925-68072) for SERCA2a for 1 hr at room temperature. The final images were detected with LiCOR Odyssey, and chemo doc and protein bands were quantified using Imagestudio-LiCOR, software-2019, and FIJI-ImageJ.

### 2.6 Single-cell isolation

Ventricular myocytes were enzymatically dissociated from young and old WT and KO mice. In short, immediately after cardiac excision, the heart was cleaned, and the aorta was cannulated. The heart was then retrogradely perfused at 2 mL/min at 36 ± 0.5°C, for 3 min with Ca^2+^ free Tyrode solution (in mM): NaCl 137, KCl 5.4, HEPES 10, MgCl_2_ 1, and glucose 10 (pH 7.3) until the effluent was clear of blood. Then the heart was perfused with the same solution containing 1 mg/mL collagenase Type A (Catalog # 9036-06-0; Roche, Germany) and 0.08 mg/mL protease Type XIV (Cat log # 48655321; Sigma-Aldrich, USA) for 8 to 11 min, followed by Tyrode solution containing 0.2 mM CaCl_2_ for 5 min. Single cells were then obtained by dissociation via gentle agitation of digested ventricular cardiac tissues. Ventricular cardiomyocyte suspensions were filtered through a nylon mesh, and cells were stored at room temperature in 0.2 mM CaCl_2_ Tyrode solution. All solutions used for dissection and perfusion were continuously bubbled with O_2_.

### 2.7 Cardiomyocyte contractility

We used the IonOptix system (Ion Optix, Milton, MA, USA) to measure sarcomere and whole-cell contractility in enzymatically isolated ventricular myocytes from the FXN-KO and FXN-WT mice. Ventricular myocytes were placed on an IonOptix chamber and onto the stage of an inverted microscope and then perfused with 37°C Tyrode solution containing 2 mM Ca^2+^. Cells were stimulated at a frequency of 1 Hz using a MyoPacer Field Stimulator (Ion Optix) connected to a pair of platinum wires placed on opposite sides of the chamber. The single cell, and sarcomere contractility analyzed using the Ionwizard software.

### 2.8 Confocal imaging

We measured the mitochondrial membrane potential (Δψ_m_) using tetramethylrhodamine methyl ester (TMRM) dye^27–30^. Left ventricular myocytes from the FXN-WT and FXN-KO were plated in an eight-well confocal chamber (LAB-TEK, DK, Denmark; Catalog # 155409,) for 20 minutes, and cells were incubated with TMRM (10 nmol/L; Invitrogen; Catalog # T668) for 20 minutes in Tyrode solution at 37°C and washed two times with Tyrode solution. After incubation with the dye, live cells were imaged for 20 min in 2 min intervals on a confocal microscope (Olympus-IX) using a 63X-oil objective. After 10 minutes of baseline recording, carbonyl cyanide 4-(trifluoromethoxy) phenylhydrazone (FCCP) (10 μmol/L) was added to both FXN-WT and FXN-KO ventricular myocytes, and fluorescence was recorded for 10 more minutes.

For the measurement of ROS, ventricular myocytes were incubated with MitoSOX Red (5 μM; Invitrogen; Catalog # M36008) for 30min at 37 °C, and cells were washed with Tyrode solution. MitoSOX dye was used as a marker for ROS measurements^31–33^. The live cells were imaged using Nikon 3i confocal microscope with a 63X water immersion objective.

Confocal settings were identical for FXN-WT and the FXN-KO myocytes. We utilized the ImageJ-FIJI (NIH, USA) software (https://fiji.sc/) to analyze the ROS fluorescence and Δψ_m_.

### 2.9 Oxygen Consumption Rate Measurements

To investigate how mitochondrial respiration was affected in FA, we measured OCR using XFp24 Cell Mito Stress Test (Catalog # B05423; Agilent Seahorse)^34, 35^. One day prior to the experiment, XFp24 cell culture plates were coated with gelatin/fibronectin at 37 °C. We enzymatically isolated ventricular myocytes from FXN-KO and FXN-WT mice. We dissociated myocytes in DMEM medium (Seahorse Bioscience), supplemented with 1 mM pyruvate, 2 mM glutamine, and 10 mM D-glucose. We plated 2500 myocytes/well into 24 well seahorse plates for 10 min. OCR was measured using the Seahorse Bioscience XF24 Extracellular Flux Analyzer (Seahorse Bioscience). Measurements were taken as the cells were incubated sequentially under four conditions: 1) at baseline; 2) oligomycin (1.5 μM) was added to inhibit ATP synthase and reduces OCR; 3) FCCP (1 μM), a mitochondrial uncoupler, was added to induce maximal respiration; and 4) Antimycin A (10 μM), a Complex III inhibitor, was added to obtain non-mitochondrial oxygen consumption background. The Seahorse software was used to extract the results and plotted them using GraphPad Prism. OCR was normalized to cell number per well.

### 2.10 Transmission Electron Microscopy (TEM)

We performed TEM as we previously described^36^. Left ventricular tissues were cut into 4 mm^3^, washed with 0.1M sodium cacodylate, and fixed in 2.5% glutaraldehyde for 48 hours at 4°C, then transferred to sodium cacodylate containing 1% osmium tetroxide for 2 hours. Each sample was dehydrated with a graded series of ethanol (35%, 50%, 70%, and 95% in water), and 100% dry acetone was used for final dehydration for 10 minutes at room temperature. Polymerized samples were sliced with a diamond knife and transferred to copper grids. Ultrathin sections were stained with lanthanum nitrate and uranyl acetate and imaged with the JEOL JEM-1400 TEM. Representative images were acquired using 25,000x and 100,000x and recorded with Digital Micrograph software.

## 3. RESULTS

### 3.1 Cardiac abnormalities in the FA mouse

We examined the ECG and echocardiographic manifestations of FA in the mouse model. Surface ECG (under 1.5% Isoflurane) was conducted in 9.5 and 12 weeks old FXN-WT and FXN-KO mice. ECG abnormalities were evident in the FXN-KO mice compared to the FXN-WT. In 9.5 weeks old animals, FXN-KO mice displayed a significant increase in QT interval duration compared to the FXN-WT (figure 1). This is consistent with clinical observations where QT prolongation and T-wave inversion have been documented in patients with FA^37–41^. In the 12 weeks old FXN-KO mice, the RR, PR, QRS, QT, and QTc intervals were significantly increased (figure 2; \**p* < 0.05 and \*\**p* < 0.01; unpaired 2-sample *t*-test) ^18, 39, 41^.

**Figure 1.**
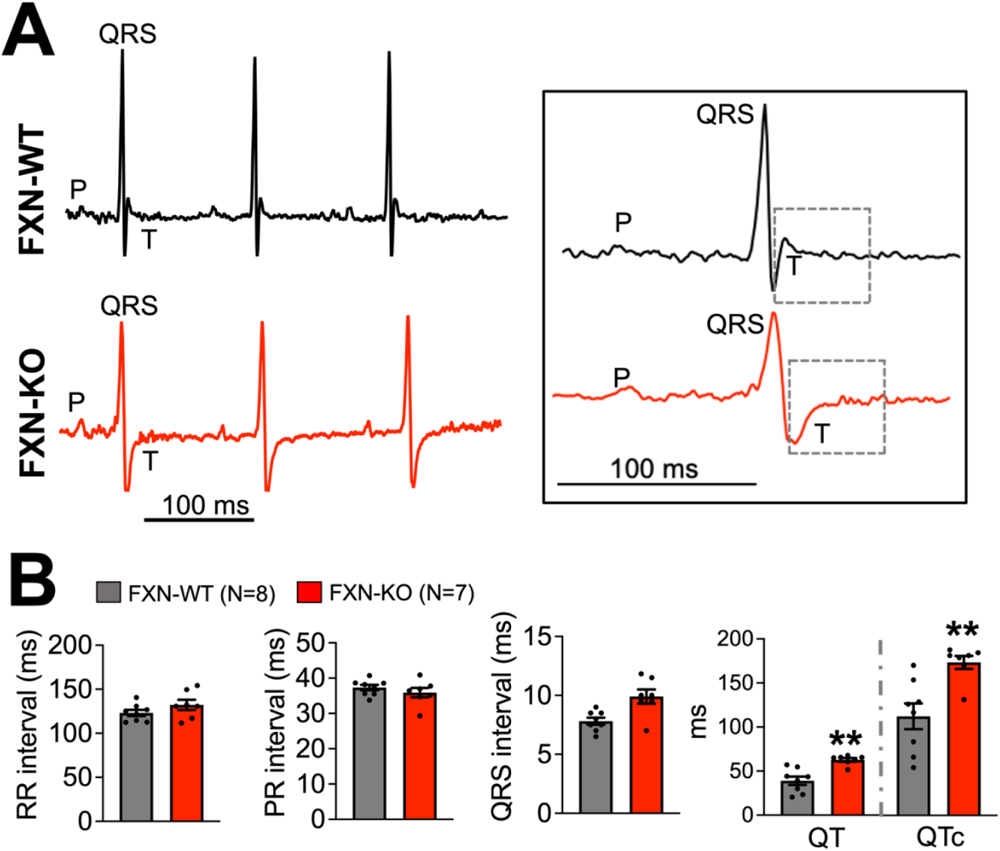
QT prolongation in 9.5 weeks old FXN-KO mice. (A) Representative ECG tracings for FXN-WT (black) and KO (red) mice. (B) Summary data comparing the duration of (a) RR, (b) PR, (c) QRS, (d) QT, and QTc intervals from WT vs. KO mice. QT and QTc were significantly increased in FXN-KO compared to the FXN-WT (** *p* < 0.01, vs. FXN-WT; unpaired 2-sample *t*-test). Bars are mean ± SEM.

**Figure 2.**
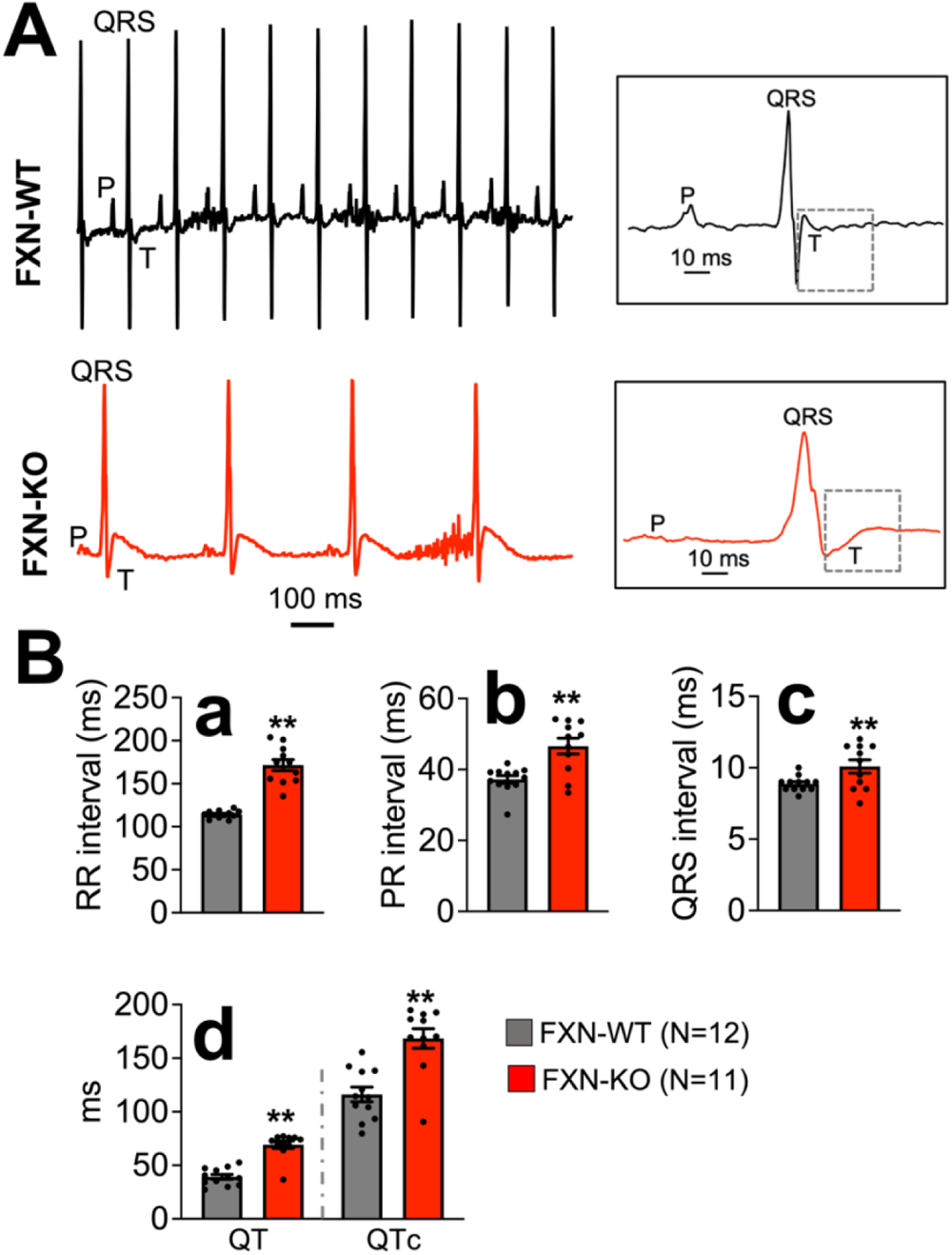
Abnormal ECG in 12 weeks old FXN-KO mice. (A) Representative ECG tracings for FXN-WT (black) and KO (red) mice. (B) Summary data comparing the duration of (a) RR, (b) PR, (c) QRS, (d) QT and QTc intervals from FXN-WT vs. KO mice. (** *p* < 0.01, vs. FXN-WT; unpaired 2-sample *t*-test). Bars are mean ± SEM.

We further assessed cardiac function and morphological developments in the FXN-KO mice using echocardiography. The M-Mode images from FXN-WT and FXN-KO suggest a significantly altered cardiac function in the FXN-KO mice (figure 3A), where the ejection fraction and fractional shortening were significantly reduced, and the left ventricular end-diastolic and end-systolic volumes were significantly increased compared to the FXN-WT (figure 3 B; \**p* < 0.05 and \*\**p* < 0.01; unpaired 2-sample *t*-test).

**Figure 3.**
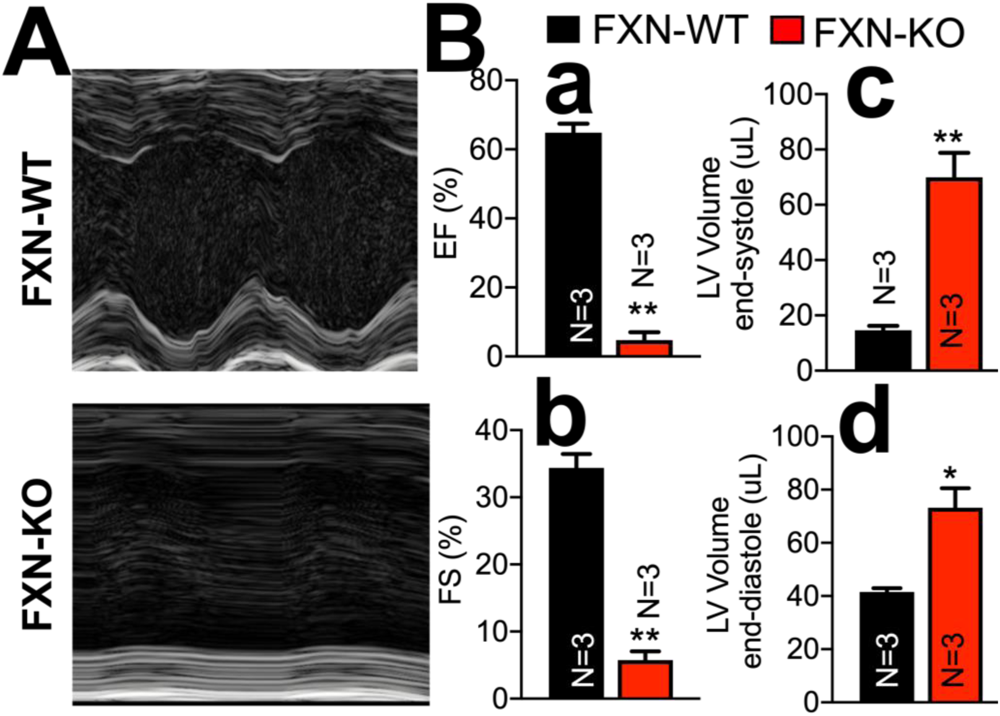
Echocardiography in FXN-WT and FXN-KO mouse hearts. B and M-mode images. (B) Summary bar graphs of (a) Left ventricular ejection fraction (EF), (b) Fractional shortening (FS), (c) Left ventricular end-systolic volume, (d) Left ventricular end-diastolic volume. (* *p* < 0.05, ** *p* < 0.01, vs. FXN-WT; unpaired 2-sample *t*-test). Bars are mean ± SEM.

These results suggest that frataxin deficiency in the mouse leads to contractile dysfunction that includes a reduced ejection fraction, similar to what has been described in FA patients^16, 42^. Altogether, this mouse FXN-KO model (MCK-Cre) seems to recapitulate several ECG and echocardiographic manifestations that have been described in FA patients with cardiac involvement.

### 3.2 Frataxin deficiency alters the major Ca^2+^ handling proteins expression in the FA heart

We conducted western blot analysis in human and mouse heart samples to possible protein abundance changes in pathways important for intracellular Ca^2+^ handling. This is since cardiac dysfunction, specifically reduced left ventricular ejection fraction, occurs in FA patients with cardiac involvement and is shown to indicate poor prognosis ^1, 43–45^. Interestingly, we found significantly decreased SERCA2a and Ryr2 expression in the human FA samples and in the FXN-KO heart samples compared to respective controls (figure 4). In figure 4, the upper panels are representative blots of SERCA2a, and Ryr2, in mouse (A) and human (C) samples. The graphs in (B) and (D) show the quantification of the SERCA2a, and Ryr2 proteins expressions which were significantly downregulated in the human and mouse FA heart samples compared to controls (\*\**p* < 0.01; unpaired 2-sample *t*-test). Overall, these results indicate that this mouse model of FA protein changes in Ca^2+^ handling, similar to the human heart with FA^19, 46, 47^.

**Figure 4.**
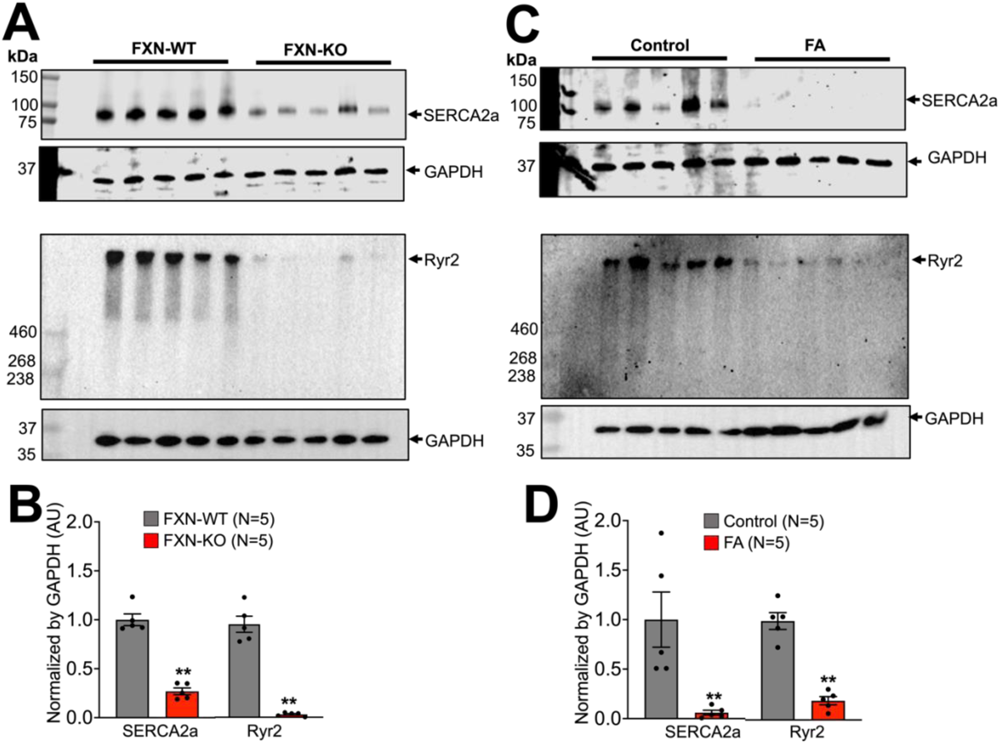
Western blot analysis of major Ca ^2+^ handling proteins levels in FA. Representative blots of sarco-endoplasmic reticulum ATPase 2a (SERCA2a) and ryanodine receptor (RyR2) in FXN-WT and KO (A), and human control and FA (C) ventricular tissue samples. Barographs are quantification of SERCA2a and RyR2 normalized to GAPDH. SERCA2a and RyR2 were significantly decreased in FXN-KO compared to the WT (B) and in human FA samples compared to the control (D). (** *p* < 0.01 vs. control or FXN-WT; unpaired 2-sample *t*-test). Bars are mean ± SEM.

### 3.3 Cellular contractile phenotype of frataxin deficiency in ventricular myocytes

We utilized the IonOptix system to measure sarcomere shortening and whole cell shortening (figure 5). The sarcomere and whole cell shortening were significantly reduced in FXN-KO myocytes compared to FXN-WT myocytes (figure 5; \**p* < 0.05; unpaired 2-sample *t*-test). We further analyzed the kinetics of sarcomere and whole cell contractility for the FXN-WT and FXN-KO ventricular myocytes (Table 2). For the sarcomere shortening, we found that the magnitude of the departure and return velocities were significantly reduced, and the time to relaxation of 90% was significantly increased in FXN-KO compared to FXN-WT myocytes (Table 2; \**p* < 0.05 and \*\**p* < 0.01; unpaired 2-sample *t*-test). For the whole cell shortening, we found that the return velocity was significantly reduced in FXN-KO compared to FXN-WT myocytes (Table 2; \**p* < 0.05; unpaired 2-sample *t*-test).

**Figure 5.**
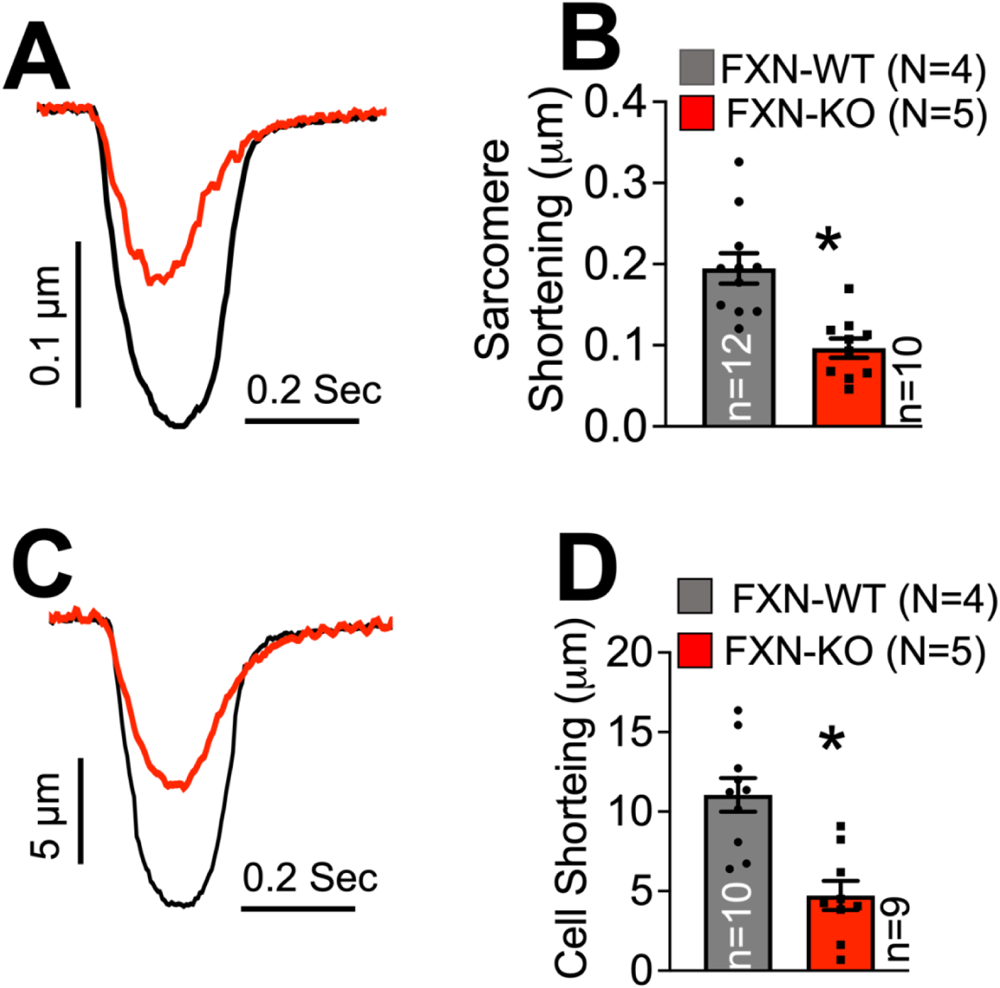
Sarcomere and cell shortening in isolated ventricular cardiomyocytes from FXN-WT and KO mice. (A) Representative traces of sarcomere shortening, (B) Summary of sarcomere shortening in FXN-KO and WT, (C) Representative traces of cell shortening, and (D) Summary of cell shortening in FXN-KO and WT. (\**p* < 0.05, vs FXN-WT; unpaired 2-sample *t*-test). Bars are mean ± SEM.

**Table 2.** Kinetics of sarcomere and whole cell contraction and relaxation in ventricular myocytes from FXN-WT and KO. Data represent mean ± SEM (\**p* < 0.05, \*\**p* < 0.01, vs. FXN-WT; unpaired 2-sample t-test).

|  | <u>**FXN-WT**</u> | <u>**FXN-KO**</u> |
| --- | --- | --- |
| <u>**Sarcomere shortening**</u> |  |  |
| Departure Velocity (µm/Sec) | -3.5 ± 0.43 | -1.77 ± 0.33 * |
| Return Velocity (µm/Sec) | 2.70 ± 0.36 | 1.10 ± 0.27 ** |
| Time to Peak 90% (Sec) | 0.87 ± 0.004 | 0.10 ± 0.006 |
| Time to Relaxation 90% (Sec) | 0.28 ± 0.01 | 0.39 ± 0.04 * |
| <u>**Whole cell shortening**</u> |  |  |
| Departure Velocity (µm/Sec) | -215.66 ± 76.82 | -94.92 ± 19.12 |
| Return Velocity (µm/Sec) | 125.41 ± 35.46 | 44.99 ± 9.01 * |
| Time to Peak 90% (Sec) | 0.092 ± 0.007 | 0.098 ± 0.01 |
| Time to Relaxation 90% (Sec) | 0.33 ± 0.05 | 0.39 ± 0.04 |

### 3.4 Lack of frataxin triggers mitochondrial perturbations and necrotic cardiac cell death in FA

The transfer of electrons via electron transport chain complexes on the inner mitochondrial membrane is essential for establishing the electromotive force and proton gradient for maintaining Δψ_m_. It is worth noting that mitochondrial energy production relies heavily on FeS-clusters (respiratory complexes I, II, and III) and heme (respiratory complex IV) as prosthetic groups, but the biogenesis of FeS-clusters itself takes place in mitochondria, which contain the specific assembly machinery. Frataxin was shown to interact with multiple core components of this machinery.

We assessed mitochondrial function by evaluating the Δψ_m_, which is a direct measurement of H^+^ ion production from proton pumps such as complex I, III, and IV in the electron transport chain during oxidative phosphorylation^27^. The Δψ_m_ was significantly depolarized in FXN-KO ventricular myocytes compared to the FXN-WT (figure 6). Figure 6 A shows representative microscopic images from FXN-WT (upper panel) and FXN-KO (lower panel). Figure 6 B is the time-dependent Δψ_m_ before and after the addition of 10 µM carbonyl cyanide 4-(trifluoromethoxy) phenylhydrazone (FCCP), a proton uncoupler that decreases and collapses the Δψ_m_. Figure 6 C shows the average change in Δψ_m_ intensity after the addition of FCCP. The average of Δψ_m_ was significantly lower in FXN-KO versus FXN-WT (\*\**p*<0.01 vs. FXN-WT, t-test, N=1; n=6).

**Figure 6.**
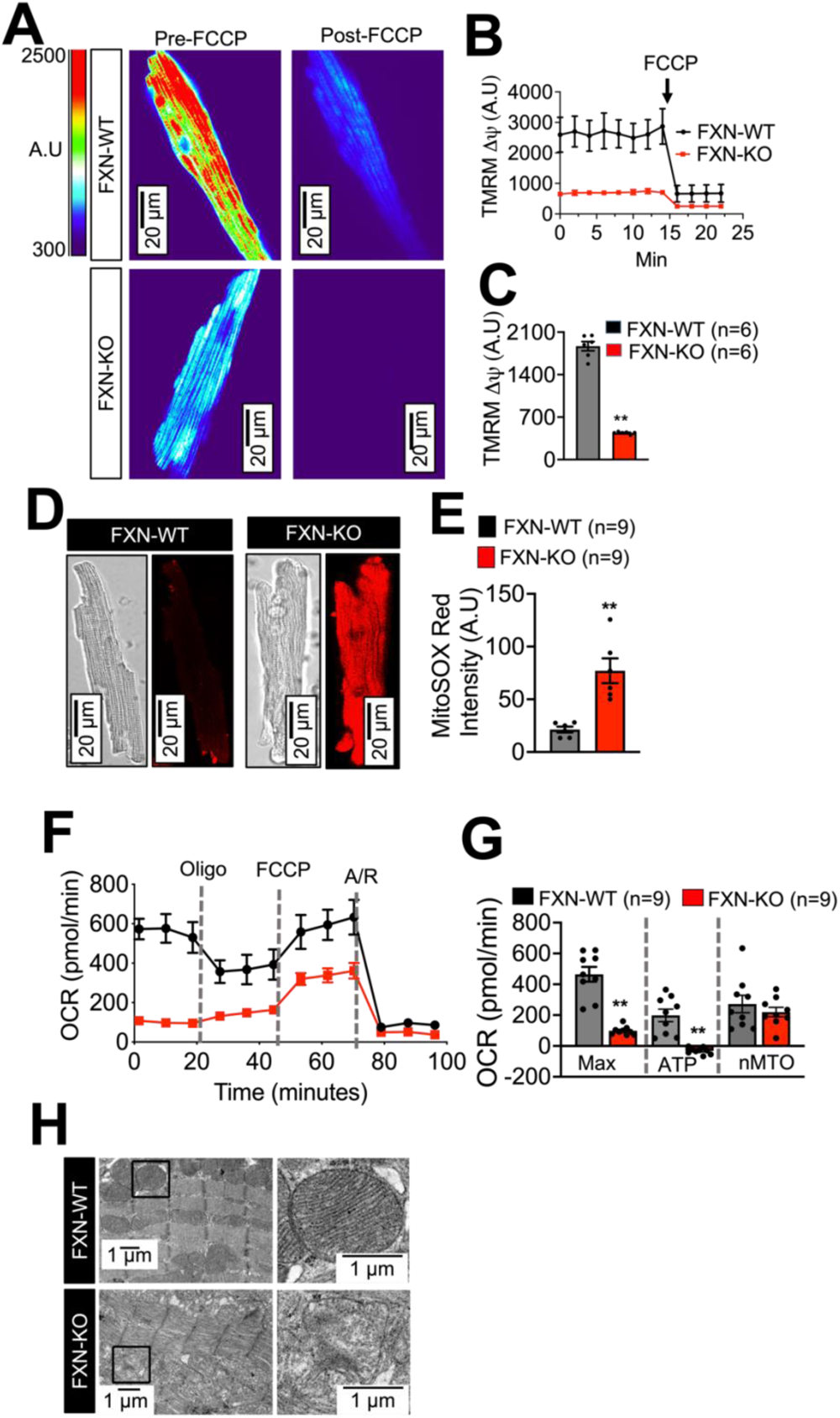
Mitochondrial perturbations in FA. (A) TMRM ΔΨm fluorescence in FXN-WT and KO cardiomyocytes. (B) Time-dependent ΔΨm over 22 min. After 15 min, the addition of 10 µM of FCCP decreased ΔΨm. (C) Average ΔΨm intensity was significantly less in FXN-KO (n=6). (D) Representative bright field and MitoSOX fluorescence images of cardiomyocytes from FXN-WT and FXN-KO. (E) Relative MitoSOX fluorescence was significantly higher in FXN-KO compared to the FXN-WT (n=9). (F) Oxygen consumption rate (OCR) in FXN-WT and FXN-KO showing the change in OCR after injections of Oligo (oligomycin), FCCP (phenylhydrazone), and A/R (Antimycin and Rotenone). (G) Quantification of normalized Max (maximal) OCR, ATP (ATP linked) OCR, and nMTO (non-mitochondrial) OCR (n=9). (H) Electron micrographs of cardiac muscle from FXN-WT and KO ventricle tissues. Left-side images were captured with 25,000X magnification, and right-side images were selected area of left-side images captured with 100,000X magnification show normal mitochondria with healthy cristae in FXN-WT and over-fused mitochondrial with disturbed cristae in FXN-KO. (** *p* < 0.01, vs. FXN-WT; unpaired 2-sample t-test). Bars are mean ± SEM.

We tested whether the observed loss of Δψ_m_ is accompanied by elevation of ROS in FA. Figures 6 D and E shows the representative fluorescence images from FXN-KO and FXN-WT. Confocal settings were identical for both conditions. Figure 6 E is the quantification of the relative fluorescence of ROS staining, which was significantly higher in FXN-KO versus control FXN-WT cells (\*\**p*<0.01 vs. FXN-WT, t-test, N=1; n=9). This experiment confirmed the greater levels of ROS in FXN-KO compared to the FXN-WT.

We measured OCR to confirm that the ROS elevation might be directly related to a disruption of respiratory chain activity. We used XFp24 Cell Mito Stress Test (Agilent Seahorse) and enzymatically isolated ventricular myocytes from FXN-KO and FXN-WT mice (figures 6 F and G). Oxygen is consumed to generate ATP in the mitochondria via the oxidative phosphorylation pathway, and therefore, measuring oxygen consumption will allow for a readout of mitochondrial respiration and cellular metabolism. After measurement of basal OCR, cellular respiration modulators, oligomycin, FCCP, and antimycin A, were injected in each well in order to quantify ATP-linked respiration and maximal respiration. Oligomycin decreased OCR by inhibiting ATP synthase. FCCP, a proton uncoupler, increased OCR to maximal levels by uncoupling oxygen consumption from ATP production. Antimycin A reduced OCR by targeting the electron transport chain^34, 35^. Figure 6 F demonstrates the change of OCR after injection of mitochondrial modulators in ventricular myocytes from the FXN-WT and FXN-KO mice, and figure 6 G shows that the basal respiratory rate, as well as ATP-linked respiration, were significantly reduced in FXN-KO myocytes (\*\**p* < 0.01, t-test). Respiratory capacity was not changed in FXN-KO mice ventricular myocytes in contrast to FXN-WT myocytes (*p* =0.21, t-test). We also assessed cardiomyocyte ultrastructure using TEM. FXN-KO mice mitochondria displayed severe alterations with sparse cristae, compared to FXN-WT mice mitochondria. Additionally, the total volume of mitochondria in FXN-KO heart tissues appears to be greater than that in FXN-WT heart tissues in figure 6 H. Thus, these results are consistent with previous findings that suggest a decreased bioenergetic efficiency and altered mitochondrial proliferation^26, 48^.

## 4. DISCUSSION

Even though FA is considered a neurodegenerative disorder, more than 50% of FA patients die from heart failure. However, the molecular mechanism that may link frataxin deficiency to cardiac dysfunctions are not well understood. In this study, we show that a loss in frataxin is associated with mitochondrial perturbations and cell death in ventricular myocytes likely through the dysregulation of Ca^2+^ handling proteins.

Our western blotting data confirmed the significant downregulation of SERCA2a, and Ryr2 in FXN-KO mice and FA human hearts compared to respective controls. Extensive research has been conducted, in animal models as well as in human subjects, to investigate the role of SERCA2a in heart failure^49, 50^. Alterations in SERCA2a expression and activity lead to a decrease in SR Ca^2+^ load and cardiac dysfunction. It has been shown that decreased SERCA2a expression may lead to cardiac hypertrophy and increased SERCA2a affinity^51^. Decreased SERCA2a has also been known to affect Ca^2+^ signaling by slowing down the rate of decay of the systolic Ca^2+^ transient, decreasing the SR Ca^2+^ content, and thus contraction^52^. Decreased Ryr2 is different than what has been described in classical heart failure, where Ryr2 was upregulated ^53–55^. Enhanced Ryr2-mediated SR Ca^2+^ leak has been observed in patients with heart failure and in various animal models^56, 57^, and the decrease of Ryr2 is unique in FA heart failure compared to classical heart failure.

It is known that the decreased activity of SERCA2a has effects on Ca^2+^ signaling. It slows down the rate of decay of the systolic Ca^2+^ transient, decreases the SR Ca^2+^ content, and thus contraction^52^. In fact, in our data, the rate of contraction (departure velocity) and relaxation (return velocity) were significantly down-regulated (Table 2) in FXN-KO compared to the FXN-WT.

In a previous study, we reported that the Δψ_m_ was depolarized, and ROS levels were elevated in human induced pluripotent stem cell-derived myocytes from a patient with FA^58^. In this study, we reconfirmed these data in ventricular myocytes from both FXN-KO and FXN-WT, and we found the Δψ_m_ was depolarized, and ROS levels were significantly elevated in FXN-KO ventricular myocytes compared to the FXN-WT. We also found significantly decreased OCR in FXN-KO ventricular myocytes compared to the FXN-WT. In previous findings, the mitochondrial structure was disorganized in FA^26, 48^. In our TEM data, we also found disrupted myofibrils and an irregular mitochondrion that were disorganized with central tubular cristae in FXN-KO ventricular myocardium compared to the FXN-WT. The increase in ROS might lead to a decrease in OCR and Δψ_m_ depolarization and structural changes in mitochondria. Importantly, it has also been shown that decreasing expression or function of SERCA2a leads to increases in oxidative stress^59^.

## 5. CONCLUSION

Treatments for cardiac dysfunctions in FA patients remain inadequate, with beta-blockers, ACE inhibitors, and diuretics being used for the management of cardiomyopathy. Additionally, oxidative stress is increased in FA, and antioxidant supplementation has been proposed as a therapy, including Vitamin E, Coenzyme Q10, and its analog Idebenone in FA patients^60, 61, 62^. These treatments do not offer protection against cardiac dysfunction progression in FA. This could be due to a lack of understanding of the EC coupling mechanism and Ca^2+^ handling in FA heart failure. In this study, we investigated the EC coupling mechanism and Ca^2+^ handling in FA hearts, and we found that major Ca^2+^ handling proteins were dysregulated in human and mouse cardiac tissue. Importantly, SERCA2a and Ryr2, which are SR Ca^2+^-release and reuptake channels involved in EC coupling, were downregulated in FA. In non-FA heart failure, Ryr2 protein expression and activity are upregulated by the phosphorylation of Ryr2 via activation of CaMKII enhanced by adrenergic activity, which can potentiate SR Ca^2+^ leak and lead to arrhythmia. In contrast, we found decreased expression of Ryr2 in FA hearts. Further investigation is needed to uncover the intracellular signaling mechanisms involved in the downregulation of major Ca^2+^ handling proteins SERCA2a in FA.

## ACKNOWLEDGEMENTS

This work was supported in part by the American Heart Association Career Developmental award to BC and the National Institutes of Health grant R01ES032099 to SN.

